# Flavor-specific enhancement of electronic cigarette liquid consumption and preference in mice

**DOI:** 10.1101/862524

**Authors:** AL Wong, SM McElroy, JM Robinson, SM Mulloy, FK El Banna, AC Harris, MG LeSage, AM Lee

**Affiliations:** Department of Pharmacology, University of Minnesota, Minneapolis, MN, USA; Graduate Program in Neuroscience, University of Minnesota, Minneapolis, MN, USA; Department of Medicine, University of Minnesota, Minneapolis, MN, USA; Department of Medicine, Hennepin Healthcare Research Institute, Minneapolis, MN, USA; Department of Psychology, University of Minnesota, Minneapolis, MN, USA

**Keywords:** Electronic cigarette, nicotine, mice, preference, aversion, consumption

## Abstract

**Background:** The use of electronic cigarettes has increased over the past decade. To determine how the abuse liability of electronic cigarette liquids (e-liquids) differs from nicotine alone, and to determine the impact of flavor, we compared nicotine-containing fruit- and tobacco-flavored e-liquids, and their nicotine-free versions, to nicotine alone in mouse models of oral consumption, reward and aversion.

**Methods:** Adult male C57BL/6J mice voluntarily consumed oral nicotine, equivalent nicotine concentrations of fruit- and tobacco-flavored e-liquid, and equivalent dilutions of the nicotine-free versions in 2-bottle choice tests. Conditioned place preference and place aversion were assessed with peripherally administered e-liquids or nicotine. Serum nicotine and cotinine levels were measured after subcutaneous injections of e-liquid or nicotine.

**Results:** Mice showed higher consumption and preference for the fruit-flavored e-liquid compared with nicotine alone. This increase was not due to the flavor itself as consumption of the nicotine-free fruit-flavored e-liquid was not elevated until the highest concentration tested. The increased consumption and preference were not observed with the tobacco-flavored e-liquid. The conditioned place preference, place aversion and nicotine pharmacokinetics of the fruit-flavored e-liquid were not significantly different from nicotine alone.

**Conclusions:** Our data suggest that fruit, but not tobacco flavor, increased the oral consumption of e-liquid compared with nicotine alone. Moreover, this enhancement was not due to increased consumption of the flavor itself, altered rewarding or aversive properties after peripheral administration, or altered pharmacokinetics. This flavor-specific enhancement suggests that some flavors may lead to higher nicotine intake and increased use of e-liquids compared with nicotine alone.

**Highlights:**

- Fruit flavor, but not tobacco flavor, enhances e-liquid consumption and preference
- The nicotine-free flavored e-liquid is not preferred over nicotine alone
- Conditioning rewarding and aversive effects are equal between nicotine and e-liquid

## 1. Introduction

Electronic cigarettes (e-cigarettes) have steadily increased in popularity over the last decade (Chou et al., 2017). Over 2 million middle and high school students have used e-cigarettes, prompting the FDA to declare e-cigarette use a youth epidemic (Gottlieb, 2018; Wang et al., 2018). Alarmingly, 33% of e-cigarette users have never used combustible cigarettes, indicating that these products are appealing to and capturing a new population that may progress to nicotine dependence (McMillen et al., 2015). Indeed, e-cigarette use is significantly associated with nicotine use disorder and nicotine addiction (Chou et al., 2017), and youth who use e-cigarettes are more likely to become combustible cigarette smokers later in life (Leventhal et al., 2015; Loukas et al., 2018).

E-cigarettes vaporize a liquid (e-liquid) that contains nicotine and flavors in a mixture of propylene glycol and glycerin. The levels of tobacco-related chemicals in e-liquids are very low due to the lack of tobacco (Han et al., 2016; Beauval et al., 2017). However, e-liquids contain other unknown chemicals and e-cigarettes can deliver as much nicotine as a combustible cigarette (Wagener et al., 2017). How the abuse liability of e-liquids differs from nicotine alone has not been extensively studied, as the majority of pre-clinical studies on e-liquids have focused on toxicity in peripheral organ systems (El Golli et al., 2016; Garcia-Arcos et al., 2016; Golli et al., 2016b; Vivarelli et al., 2019). The neurocognitive effects and addiction-relevant properties of e-liquids are beginning to be examined in rodent models (Golli et al., 2016a; LeSage et al., 2016a; LeSage et al., 2016b; Harris et al., 2017; Harris et al., 2018b; Smethells et al., 2018). Intriguingly, two of these studies suggest that high concentrations of e-liquids are less aversive compared with equivalent concentrations of nicotine alone in a model of intra-cranial self-stimulation (ICSS) in male rats (LeSage et al., 2016b; Harris et al., 2018b). In addition, e-liquids are available in many different flavors, and some of the most popular flavors among youth are mint, mango and fruit (Leventhal et al., 2019). The impact of different flavors is only beginning to be determined in preclinical models, with the majority of studies focusing on menthol (Alsharari et al., 2015; Wickham, 2015; Henderson et al., 2019).

In this study, we compared the voluntary consumption and preference of fruit- and tobacco-flavored e-liquids to nicotine alone in a two-bottle choice model in mice. Two-bottle choice is a high-throughput, technically simple assay that is commonly used to measure the voluntary oral consumption and preference of nicotine in mice (Klein et al., 2004; Glatt et al., 2009; Lee and Messing, 2011; Cao et al., 2012; Locklear et al., 2012; O’Rourke et al., 2016). Although the pharmacokinetics of oral consumption are slower compared with intravenous nicotine delivery, voluntary nicotine consumption in mice can lead to physical dependence and is regulated by the same genetic and molecular factors that modulate nicotine intake in humans, such as enzymatic regulation of nicotine metabolism and expression of nicotinic acetylcholine receptors (nAChRs) (Siu et al., 2006; Locklear et al., 2012; Renda et al., 2016; Bagdas et al., 2019).

We found that mice showed greater consumption and preference for fruit-flavored e-liquid, but not tobacco-flavored e-liquid, compared with equivalent concentrations of nicotine alone. This increase was not due to the flavor itself, as consumption and preference of a nicotine-free fruit-flavored e-liquid was not elevated until the highest concentration tested. We then assessed whether fruit-flavored e-liquid had altered rewarding or aversive properties compared with nicotine alone in the conditioned place preference (CPP) and conditioned place aversion (CPA) assays, and found no significant differences compared with nicotine alone. Our data suggest that fruit, but not tobacco flavor, acts to enhance oral nicotine consumption in mice. This suggests that some flavors may lead to higher nicotine intake and result in altered abuse liability of e-liquids compared with nicotine alone.

## 2. Methods

### 2.1. Animals and reagents

Eight-week old male C57BL/6J mice from The Jackson Laboratory (Sacramento, CA) acclimated to our facility for at least one week before behavioral experiments. Mice were group housed in standard cages under a 12-h light/dark cycle until the start of experiments, after which they were individually housed. All animal procedures were in accordance with the Institutional Animal Care and Use Committee at the University of Minnesota, and conformed to NIH guidelines.

Nicotine tartrate salt (Acros Organics, Thermo Fisher Scientific, Chicago, IL) was mixed with tap water to the concentrations reported for each experiment. The e-liquids Retro Fruit Twist and Classic American Tobacco were purchased from NicVape.com, and consisted of a 50/50 propylene glycol and glycerin mix. All e-liquid solutions were verified for their nicotine content by a standard gas chromatography assay with nitrogen phosphorus detection, based on the method of Jacob and colleagues (Jacob et al., 1981; Hieda et al., 1999; LeSage et al., 2003; Harris et al., 2008). The actual nicotine concentrations in the fruit- and tobacco-flavored e-liquids were between 16.1 to 17.7 mg/mL, and the nicotine content of the nicotine-free e-liquids were between 0.000123 to 0.000655 mg/mL (labelled nicotine concentrations=18 and 0 mg/mL). All concentrations were reported as free base. The nicotine and e-liquid solutions for voluntary consumption experiments were diluted in tap water, and the solutions for peripheral injections were pH adjusted to 7.4 and diluted in 0.9% saline.

### 2.2. Voluntary oral drug consumption (2-bottle choice tests)

Two-bottle choice consumption was performed in a similar manner as our prior work (O’Rourke et al., 2016; Touchette et al., 2018; DeBaker et al., 2019). For each group, the mice were singly housed and presented with one bottle of tap water and one bottle of drug formulation diluted in tap water (either nicotine alone, fruit-flavored nicotine-containing e-liquid, tobacco-flavored nicotine-containing e-liquid, nicotine-free fruit-flavored e-liquid, or nicotine-free tobacco-flavored e-liquid). The concentrations presented were 30, 50, 75, 100 and 200 μg/mL nicotine, with each concentration presented for one week. The nicotine-containing e-liquids were diluted to the desired nicotine concentrations, and the nicotine-free e-liquids were diluted with the same volume of water to match the nicotine-containing e-liquids. The bottles were weighed every 2-3 days and the positions of the bottles were alternated each weighing to control for side preferences. All solutions were refreshed every 3-4 days. The mice were weighed once a week, and food was freely available at all times.

### 2.3. Place conditioning

We used 0.5 and 2.0 mg/kg nicotine, or equivalent concentrations of nicotine-containing e-liquid, for CPP and CPA, respectively. To determine whether nicotine-free fruit-flavored e-liquid had any effects alone, we compared it to saline. The chamber apparatus consisted of a two-compartment place preference insert in an open field chamber with different floor textures (Med Associates, St. Albans, VT). We used an unbiased nicotine place conditioning procedure as previously reported (Grabus et al., 2006; Lee and Messing, 2011), which consisted of one habituation session on Day 1, twice daily conditioning sessions on Days 2-4, and one test session on Day 5. For the habituation session, mice were *i.p.* injected with saline and placed in the apparatus with access to both chambers for 15 minutes. For the conditioning sessions, mice were *i.p.* injected with the drug formulation and were immediately confined to one chamber for 30 minutes. Four to five hours later, mice received an injection of saline paired with the alternate chamber, and this was repeated for 3 days for a total of 6 conditioning sessions (3 drug formulation and 3 saline). On test day, mice received an injection of saline and access to both chambers for 15 minutes. The order of the injections and the drug formulation-paired floor was counterbalanced across groups. Saline control mice received saline paired with both floors. The experiments for each formulation and concentration were performed in multiple cohorts over several months.

### 2.4. Nicotine and cotinine pharmacokinetics

Mice were subcutaneously injected with 2.5 mg/kg nicotine or equivalent nicotine concentrations of e-liquid and sacrificed at 10, 20, 30 or 50 minutes after injection. Trunk blood was collected for assessment of serum nicotine and cotinine concentrations as described previously (Jacob et al., 1981; Hieda et al., 1999; LeSage et al., 2003; Harris et al., 2008).

### 2.5. Statistical analysis

For the oral consumption experiments, we calculated nicotine consumption (mg/kg) and preference for the drug formulation bottle. The consumption (mg/kg) was calculated based on of the weight of the fluid consumed and mouse weights. For the nicotine-free e-liquids, the consumption is calculated as a hypothetical mg/kg to compare with the e-liquid and nicotine groups. The preference was calculated as the weight of fluid consumed from the drug formulation bottle divided by the total fluid consumed multiplied by 100. For the place conditioning experiments, we calculated a conditioning index, which was the time spent in the drug-paired chamber during test day minus time spent in that same chamber on habituation day. All analyses were calculated using Prism 8.0 (GraphPad, La Jolla, CA). For the place conditioning data, outliers were identified using the Grubb’s test or if the data point was outside 2X the standard deviation from the mean. The number of outliers per group were: 2 for saline, 1 each for the nicotine-free e-liquid, 0.5 mg/kg nicotine, 0.5 mg/kg e-liquid, and 2.0 mg/kg nicotine groups. The determination of whether place conditioning produced preference or aversion was established by analyzing data using one-sample *t*-tests against a hypothetical conditioning index of zero for each group. Drug formulation groups were compared using Student’s *t*-tests or one-way ANOVA followed by Tukey’s multiple comparisons tests. Comparison of data across time used two-way repeated measures ANOVA followed by Tukey’s multiple comparisons tests.

## 3. Results

### 3.1. The consumption and preference of fruit-flavored e-liquid compared with nicotine alone

We compared the average daily consumption of the nicotine-containing fruit-flavored e-liquid and equivalent dilutions of the nicotine-free fruit-flavored e-liquid to nicotine alone. We found a significant interaction between drug formulation and concentration (F_interaction_(8, 148)=14.20, *P*<0.0001; F_concentration_(4, 148)=68.39, *P*<0.0001; F_drug_(2, 37)=7.785, *P*=0.002; Fig. 1A). Tukey’s multiple comparisons showed that mice consumed more nicotine-containing fruit-flavored e-liquid compared with nicotine alone at the 75, 100 and 200 μg/mL concentrations. The consumption of nicotine-containing fruit-flavored e-liquid was also significantly higher than the nicotine-free version at the 75 μg/mL concentration. The consumption of the nicotine-free fruit-flavored e-liquid was greater than both the nicotine-containing fruit-flavored e-liquid and nicotine alone at only the 200 μg/mL concentration.

**Fig. 1.**
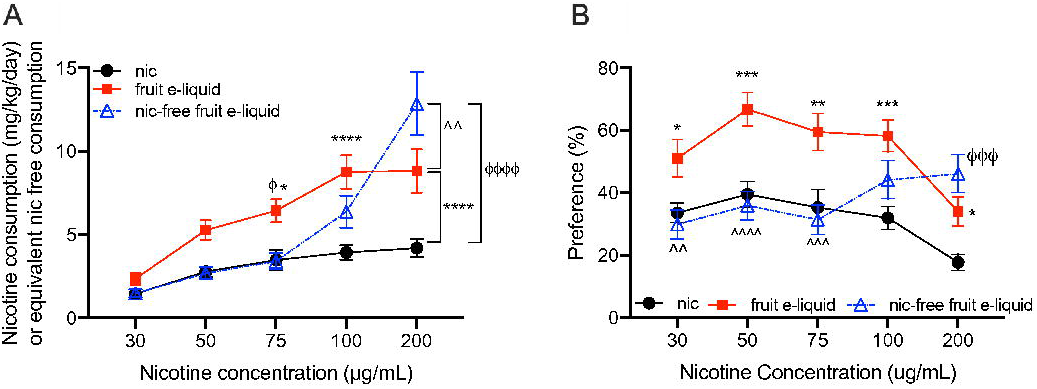
Consumption and preference of fruit-flavored e-liquid, nicotine-free fruit-flavored e-liquid and nicotine alone in 2-bottle choice tests. **(A)** The average consumption and **(B)** preference for nicotine alone, fruit-flavored e-liquid at equivalent nicotine concentrations, and nicotine-free fruit-flavored e-liquid at equivalent dilutions. The consumption for nicotine and nicotine-containing e-liquids are in mg/kg/day, and in hypothetical mg/kg/day for the nicotine-free e-liquid. \**P*<0.05, \*\**P*<0.01, \*\*\**P*<0.001 and \*\*\**P*<0.0001 for all comparisons. *indicates comparisons between fruit-flavored e-liquid and nicotine alone, ^ϕ^indicates comparisons between nicotine-free fruit-flavored e-liquid and nicotine alone, and ^^^indicates comparisons between fruit-flavored e-liquid and the nicotine-free version. Mean ± SEM, *n*=15 for nicotine alone, *n*=12 for fruit-flavored e-liquid, *n*=13 for the nicotine-free fruit-flavored e-liquid.

Similar results were observed for the bottle preference, where we found a significant interaction between drug formulation and concentration (F_interaction_(8,148)=9.820, *P*<0.0001; F_concentration_(4, 148)=11.50, *P*<0.0001; F_drug_(2, 37)=7.811, *P*=0.002; Fig. 1B). Tukey’s multiple comparisons showed that mice had greater preference for nicotine-containing fruit-flavored e-liquid compared with nicotine alone at all concentrations. There was greater preference for the nicotine-containing fruit-flavored e-liquid compared with the nicotine-free version at the 30, 50 and 75 μg/mL concentrations. The preference for the nicotine-free fruit-flavored e-liquid exceeded that of nicotine alone at the 200 μg/mL concentration. Together, these data indicate that mice increased the consumption and preference of a nicotine-containing fruit-flavored e-liquid compared with nicotine alone, and this increase is not due to increased preference for the flavor itself since consumption of the nicotine-free fruit-flavored e-liquid was not elevated until the highest concentration tested.

### 3.2. The consumption and preference of tobacco-flavored e-liquid compared with nicotine alone

The increased consumption and preference for the fruit-flavored e-liquid did not occur for the tobacco-flavored e-liquid compared with nicotine alone. For the average daily consumption, we found a significant interaction between drug formulation and concentration (F_interaction_(8, 160)=16.22, *P*<0.0001; F_concentration_(4, 160)=93.81, *P*<0.0001; F_drug_(2, 40)=4.591, *P*=0.02; Fig. 2A). Tukey’s multiple comparisons showed that the consumption of the nicotine-free tobacco-flavored e-liquid was significantly higher than both the nicotine-containing tobacco-flavored e-liquid and nicotine alone only at the 200 μg/mL concentration. No other significant differences between drug formulation were observed at any concentration. For bottle preference, we also found a significant interaction between drug formulation and concentration (F_interaction_(8, 160)=6.933, *P*<0.0001; F_concentration_(4, 160)=5.572, *P*=0.0003; F_drug_(2, 40)=3.103, *P*=0.06; Fig. 2B). Tukey’s multiple comparisons showed that the preference for the nicotine-free tobacco-flavored e-liquid was greater than nicotine alone at 100 μg/mL, and greater than both the nicotine-containing tobacco-flavored e-liquid and nicotine alone at the 200 μg/mL concentration. No significant differences were observed between nicotine alone and the nicotine-containing tobacco-flavored e-liquid at any concentration.

**Fig. 2.**
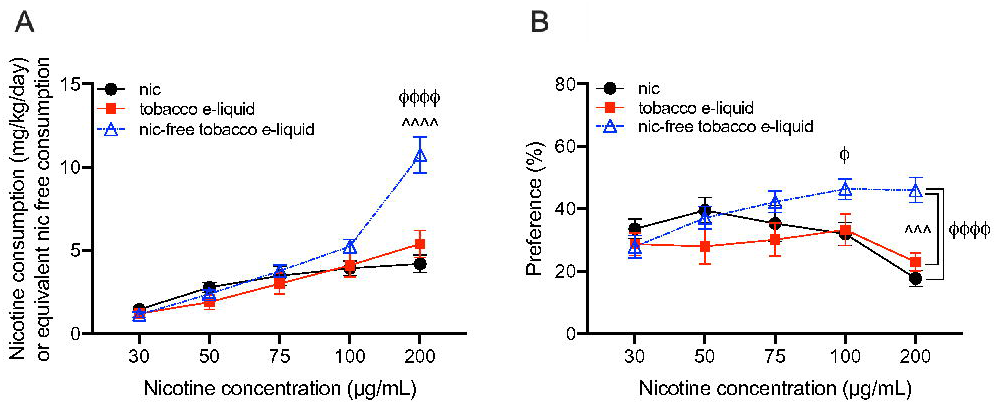
Consumption and preference of tobacco-flavored e-liquid, nicotine-free tobacco-flavored e-liquid and nicotine alone in 2-bottle choice tests. **(A)** The average consumption and **(B)** preference for nicotine alone, tobacco-flavored e-liquid at equivalent nicotine concentrations, and nicotine-free tobacco-flavored e-liquid at equivalent dilutions. The consumption for nicotine and nicotine-containing e-liquids are in mg/kg/day, and in hypothetical mg/kg/day for the nicotine-free e-liquid. \**P*<0.05, \*\**P*<0.01, \*\*\**P*<0.001 and \*\*\**P*<0.0001 for all comparisons. ^ϕ^indicates comparisons between nicotine-free tobacco-flavored e-liquid and nicotine alone, and ^^^indicates comparisons between tobacco-flavored e-liquid and the nicotine-free version. Mean ± SEM, *n*=15 for nicotine alone, *n*=14 for tobacco-flavored e-liquid, *n*=14 for the nicotine-free tobacco-flavored e-liquid.

### 3.3. The consumption and preference of nicotine-free fruit- versus nicotine-free tobacco-flavored e-liquid

We then evaluated the consumption and preference of the nicotine-free fruit-flavored e-liquid compared with the nicotine-free tobacco-flavored e-liquid to determine whether the flavors showed similar consumption and preference. For both consumption and preference, we found a main effect of concentration with no main effect of e-liquid or an interaction between e-liquid and dilution (mg/kg/day consumption: F_interaction_(4, 100)=1.153, *P*=0.34; F_dilution_(4, 100)=88.47, *P*<0.0001; F_e-liquid_(1, 25)=0.779, *P*=0.39; bottle preference: F_interaction_(4, 100)=1.616, *P*=0.18; F_dilution_(4, 100)=13.18, *P*<0.0001; F_e-liquid_(1, 25)=0.214, *P*=0.65).

### 3.4. Place conditioning of fruit-flavored e-liquid compared with nicotine alone

We first assessed CPP using 0.5 mg/kg nicotine and equivalent concentrations of fruit-flavored e-liquid (Fig. 3A). We found no significant difference in the conditioning index between fruit-flavored e-liquid and nicotine alone (*t*=1.342, df=30, *P*=0.19). However, when assessing whether each group had significant place conditioning, we found that conditioning with 0.5 mg/kg nicotine produced a conditioning index that was significantly greater than zero, indicating a significant place preference (one-sample *t*-test: *t*=2.172, df=16, *P*=0.045). The conditioning index of the fruit-flavored e-liquid was not significantly different from zero, indicating no preference or aversion was produced (one-sample t-test: *t*=0.432, df=14, *P*=0.67).

**Fig. 3.**
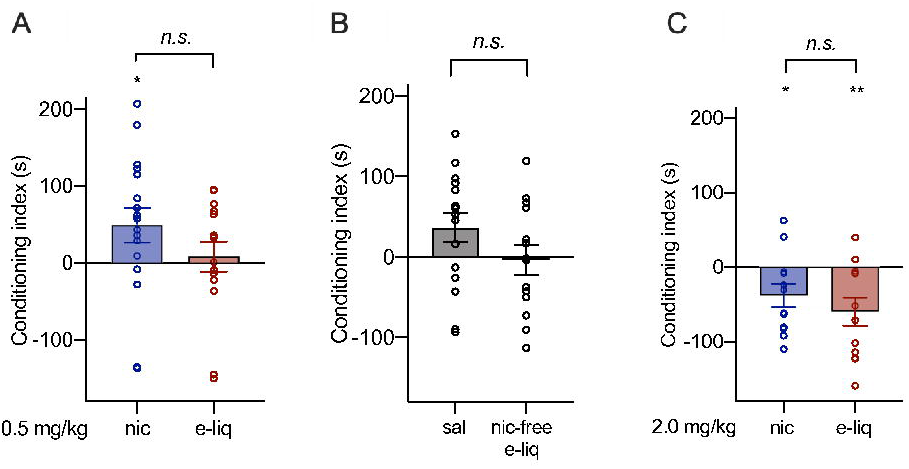
Fruit-flavored e-liquid does not differ from nicotine alone in conditioned place preference or conditioned place aversion assays. **(A)** The conditioning index after CPP with 0.5 mg/kg nicotine or fruit-flavored e-liquid at an equivalent nicotine concentration. *n*=17 for nicotine, *n*=15 for e-liquid groups. **(B)** The conditioning index after saline or nicotine-free fruit-flavored e-liquid at an equivalent dilution to 0.5 mg/kg nicotine. *n*=16 for saline, *n*=14 for nicotine-free e-liquid groups. **(C)** The conditioning index after CPA with 2.0 mg/kg nicotine or fruit-flavored e-liquid at an equivalent nicotine concentration. *n*=12 for nicotine and e-liquid groups. \**P*<0.05, \*\**P*<0.01 for a one-sample t-test between the conditioning index and a hypothetical index of zero.

We also assessed whether the nicotine-free fruit-flavored e-liquid at an equivalent 0.5 mg/kg nicotine dilution produced any preference or aversion in the place conditioning assay compared with saline alone. We found no significant difference between place conditioning with saline compared with the nicotine-free fruit-flavored e-liquid (*t*=1.553, df=28, *P*=0.13, Fig. 3B). Neither saline nor the nicotine-free fruit-flavored e-liquid produced a conditioning index that was significantly different from zero, indicating no preference or aversion was produced with either substance (one-sample t-tests saline: *t*=2.014, df=15, *P*=0.06; nicotine-free e-liquid: *t*=0.204, df=13, *P*=0.84).

We next assessed place aversion using CPA at a concentration of 2.0 mg/kg nicotine and equivalent nicotine concentrations of fruit-flavored e-liquid. There was no significant difference in the conditioning index between substances (*t*=0.896, df=22, *P*=0.38, Fig. 3C). Both nicotine alone and the fruit-flavored e-liquid produced a conditioning index that was significantly below zero, indicating that both substances produced place aversion (one-sample t-tests nicotine: *t*=2.454, df=11, *P*=0.03; fruit-flavored e-liquid: *t*=3.196, df=11, *P*=0.009). Overall, these data show that *i.p.* administered fruit-flavored e-liquid produced similar effects compared with nicotine alone, suggesting that the enhancement in fruit-flavored e-liquid consumption and preference was not due to altered reward or aversion when the drugs are administered peripherally.

### 3.5. Nicotine and cotinine pharmacokinetics

We assessed the pharmacokinetics of nicotine and cotinine after a 2.5 mg/kg *s.c.* injection of nicotine alone, and equivalent concentrations of fruit-flavored and tobacco-flavored e-liquids. For the average serum nicotine levels (ng/mL), we found a significant interaction between drug formulation and time (F_interaction_(6, 20)=2.925, *P*=0.03; F_time_(3, 20)=95.03, *P*<0.0001; F_drug_(2, 20)=35.56, *P*<0.0001; Fig. 4A). Tukey’s multiple comparisons showed that serum nicotine levels after the injection of tobacco-flavored e-liquid was significantly higher than nicotine alone at the 30 and 50 minute timepoints, and significantly higher than the nicotine-containing fruit-flavored e-liquid at the 10, 30 and 50 minute timepoints. There was no difference between the fruit-flavored e-liquid and nicotine alone at any timepoint.

**Fig. 4.**
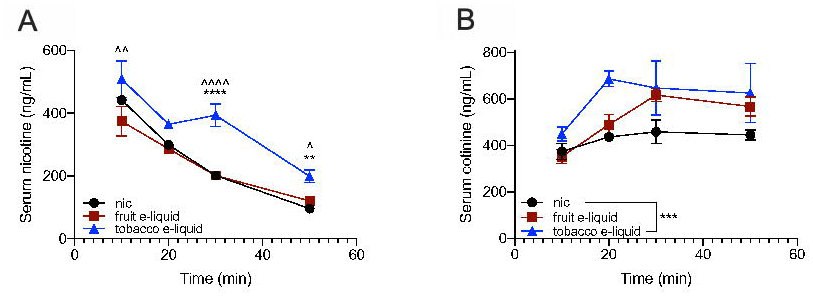
Nicotine and cotinine clearance. **(A)** The average serum nicotine and **(B)** serum cotinine levels after injection of 2.5 mg/kg *s.c.* of nicotine alone, nicotine-containing fruit-flavored e-liquid and nicotine-containing tobacco-flavored e-liquid. \**P*<0.05, \*\**P*<0.01, \*\*\**P*<0.001 and \*\*\**P*<0.0001 for all comparisons. *indicates comparison between tobacco-flavored e-liquid and nicotine alone, and ^^^indicates comparison between tobacco-flavored e-liquid and fruit-flavored e-liquid.

For the average serum cotinine levels (ng/mL), we found main effects of time and drug formulation without a significant interaction (F_interaction_(6, 24)=0.8097, *P*=0.57; F_time_(3, 24)=5.769, *P*=0.004; F_drug_(2, 24)=8.713, *P*=0.001; Fig. 4B). We examined the main effects of drug formulation using a Tukey’s multiple comparisons test and found an overall significant difference between the nicotine-containing tobacco-flavored e-liquid and nicotine alone.

## 4. Discussion

The prevalence of e-cigarette use has steadily increased over the past decade, and the numerous flavors available contributes to the popularity of these products among adolescents and young adults (McMillen et al., 2015; Leventhal et al., 2019). How the abuse liability of e-liquid differs from nicotine alone has not been extensively studied in pre-clinical models, and the studies that have been published have mainly used peripheral administration routes. In this study, we evaluated voluntary oral consumption, CPP, CPA, and nicotine pharmacokinetics of fruit- and/or tobacco-flavored e-liquids compared with nicotine alone. We found that mice had greater consumption and preference for the nicotine-containing fruit-flavored e-liquid compared with nicotine alone. Interestingly, this was not due to the flavor itself, since the consumption and preference of the nicotine-free fruit-flavored e-liquid was not elevated until the highest concentration tested. One possible mechanism may be that fruit flavoring acts as an orosensory cue to enhance the reinforcing effects of moderate nicotine concentrations, similar to how light and tone cues enhance the responding to *i.v.* nicotine self-administration in rats (Chaudhri et al., 2005; Chaudhri et al., 2006). Human data shows that young adult smokers rate green apple and chocolate flavored e-cigarettes as more rewarding compared with unflavored e-cigarettes, and are willing to work harder for the flavored e-cigarettes compared with unflavored e-cigarettes (Audrain-McGovern et al., 2016). Interestingly, we did not observe an increase in consumption and preference with tobacco-flavored e-liquids, which suggests that some flavors, but not others, can enhance consumption. We believe the mice perceived the fruit and tobacco flavoring equally, as there was no difference in the consumption or preference of the nicotine-free versions of both e-liquids. Alternatively, fruit flavor, but not tobacco flavor, may mask the aversive orosensory effects of nicotine, thus promoting greater consumption. Further research on individual flavors and flavor categories will be important in identifying the nature of this enhancement of nicotine consumption and preference.

We compared the fruit-flavored e-liquid to nicotine alone in place conditioning assays to determine whether the enhancement was due to altered rewarding or aversive properties of the e-liquid. We used peripheral administration in the CPP and CPA tests, which would eliminate the taste, but perhaps not the smell, of the e-liquid. We found no significant difference in the CPP generated by 0.5 mg/kg nicotine compared with equivalent nicotine concentrations of the fruit-flavored e-liquid. We only tested 0.5 mg/kg nicotine, which is a concentration that we and others have used to successfully produce CPP in mice (Grabus et al., 2006; Lee and Messing, 2011). It is possible that the fruit-flavored e-liquid may show differences in conditioned reward compared with nicotine alone at other concentrations. We found no significant differences in the CPA produced by fruit-flavored e-liquid and nicotine alone at 2.0 mg/kg, suggesting that the aversive properties are similar.

We also did not observe any significant differences in nicotine or cotinine pharmacokinetics between the fruit-flavored e-liquid and nicotine alone after a 2.5 mg/kg *s.c.* injection, suggesting that the enhancement of oral consumption and preference is not due to altered drug clearance, which can influence nicotine intake in humans and animals (Rao et al., 2000; Siu et al., 2006). Interestingly, we found that the nicotine-containing tobacco-flavored e-liquid resulted in higher serum nicotine and cotinine levels compared with both the fruit-flavored e-liquid and nicotine alone. However, the increases in serum nicotine and cotinine levels were not associated with altered oral consumption and preference compared with nicotine alone. The mechanism underlying the higher serum concentrations is unclear, and it is possible that the fruit- and tobacco-flavored e-liquids differ in beta-nicotyrine levels, which is formed by the oxidation of nicotine and can inhibit cytochrome P450 2A enzymes, thus inhibiting nicotine pharmacokinetics (Abramovitz et al., 2015).

Together, our data suggest that the increase in consumption and preference of the fruit-flavored e-liquid is primarily due to the orosensory properties of the flavor, and not to an interaction with nicotine when administered peripherally. This has important implications for behavioral assays that require peripheral administration, such as intravenous self-administration, which may be unable to detect the orosensory effects of flavored e-liquids.

The abuse liability of e-liquids compared with nicotine alone has been understudied compared with the rapid increase in popularity of these products. Two previous studies in adult male rats using peripheral administration of a fruit-flavored e-liquid showed that high concentrations of the e-liquid are less aversive compared with nicotine alone in an ICSS model of aversion, whereas no differences were observed in an ICSS model of reward, *i.v.* self-administration, or nicotine pharmacokinetics (LeSage et al., 2016b; Harris et al., 2018b). Further investigation found that propylene glycol, a main component of all e-liquids, is able to attenuate the aversive effect of nicotine alone in the ICSS procedure, without affecting ICSS thresholds itself (Harris et al., 2018a). In this study, peripheral administration of nicotine-containing fruit-flavored e-liquid was not different from nicotine alone in the CPA procedure, indicating equal aversive conditioning was produced. This difference in findings may be due to several factors, including a species difference, differences in the doses used, in the behavioral assay, or batch differences in the composition of the fruit-flavored e-liquids. We did not assess the effect of propylene glycol in the place conditioning assay, and there is no data on how propylene glycol affects place conditioning in mice. However, we tested the effect of the nicotine-free fruit-flavored e-liquid, which contains a 50/50 propylene glycol and glycerin mixture, and did not observe any conditioned preference or aversion.

The exact chemical composition of the compounds used as flavorings in e-liquids is unknown as manufacturers are not required to provide a list of ingredients. Recent evaluation of flavor preferences of JUUL e-liquid in US youth from 8^th^ to 12^th^ grade shows that mint, mango and fruit are the most preferred (Leventhal et al., 2019). Menthol, the compound primarily used in mint flavoring, has been extensively studied, as it is the only flavor allowed in combustible cigarettes (Wickham, 2015). Menthol acts through several mechanisms, such as reduction of the aversive sensory effects of smoking in humans, attenuation of the aversion to high nicotine concentrations in two-bottle choice tests in rats (Wickham, 2015; Wickham et al., 2018), and delaying the clearance of nicotine (Alsharari et al., 2015). Unlike menthol, mango flavoring in e-liquid appears to be a combination of at least 7 chemical compounds (Eddingsaas et al., 2018). The chemical composition of fruit and tobacco flavoring in the e-liquids used in the present study is unknown, but it is highly likely that they are composed of multiple chemicals, similar to mango flavoring. Identifying whether these individual chemicals are important in the abuse liability of e-liquids will be a challenge.

In this study, one limitation is that we assessed adult male mice only. Determining whether the enhancement of fruit-flavored e-liquid consumption and preference also occurs in adult female mice and adolescent mice of both sexes is critical to understanding the biological impact of these products. In addition, the technology to enable voluntary self-administration of inhaled e-liquids is still under development. Future replication of these flavor effects in an inhalation model or through the use of aerosolized e-liquid extracts would be important to more closely model human intake.

## 5. Conclusions

We found that mice had higher consumption and preference of a fruit-flavored e-liquid compared with nicotine alone. Importantly, this was not due to the flavor itself, as the nicotine-free fruit-flavored e-liquid was not preferred until the highest concentration. Moreover, this increase in consumption and preference was not observed with the tobacco-flavored e-liquid. There was no significant difference in the CPP or CPA of the fruit-flavored e-liquid compared with nicotine, thus the increased consumption and preference were likely not due to altered rewarding or aversive effects of the e-liquid when administered peripherally. Together, our results suggest that certain flavors may enhance nicotine consumption. Identifying which flavors produce this effect, the chemical composition of the flavors, and the mechanism of the enhancement will be important in determining how the abuse liability of e-liquids may differ compared with nicotine alone, and which regulatory steps may be required to limit the abuse of these products.

## Contributors

ALW and SM McElroy collected and analyzed the voluntary consumption data, JMR performed the place conditioning experiments, SM Mulloy and FKE performed the pharmacokinetic study. MGL and ACH provided substantial assistance with the data analysis, interpretation and manuscript preparation. AML was responsible for the overall design and execution of the project, data collection, analyses and prepared the first draft of the manuscript. All authors contributed to and have approved the final manuscript.

## Acknowledgments

We thank Theresa Harmon for assistance with data collection.

